# Declining population sizes and loss of genetic diversity in commercial fishes: a simple method for a first diagnostic

**DOI:** 10.1101/2021.12.16.472909

**Authors:** Natalia Petit-Marty, Liu Min, Iris Ziying Tan, Arthur Chung, Bàrbara Terrasa, Beatriz Guijarro, Francesc Ordines, Sergio Ramírez-Amaro, Enric Massutí, Celia Schunter

**Affiliations:** School of Biological Sciences and Swire Institute of Marine Science. The University of Hong Kong, Hong Kong (SAR); State Key Laboratory of Marine Environmental Science and College of Ocean and Earth Sciences, Xiamen University, Xiamen, Fujian, P.R. China; Laboratori de Genètica, Universitat de les Illes Balears, Palma, Spain; Instituto Español de Oceanografía, Centre Oceanogràfic de les Balears, Palma, Spain

**Keywords:** adaptive potential, COI barcode, conservation, fisheries, global change, overfishing

## Abstract

Exploited fish species may have or are experiencing declines in population sizes coupled with a decrease in genetic diversity. This can lead to the loss of adaptive potential to face current and future environmental changes. However, little is known about this subject while research on it is urgently needed. Thus, this study aims to answer a simple, even naive question, given the complexity of the subject: Could we use a simple method to obtain information on the loss of genetic diversity in exploited fish species? We investigated the use of the levels of genetic diversity in the widely used genetic marker Cytochrome C Oxidase subunit I (COI) mitochondrial gene. Estimates of genetic diversity in COI were obtained for populations of seven fish species with different commercial importance from the East China Sea. These estimates were contrasted against a large dataset of fish species distributed worldwide (N=1426), a dataset of East-Asian fish species (N=118), two farmed species with expected low genetic diversity, and four long-term managed species from the Mediterranean Sea. We found that estimates of genetic diversity in COI match the expectations from theoretical predictions, known population declines, and fishing pressures. Thus, the answer to our question is affirmative and we conclude that estimates of genetic diversity in COI provide an effective first diagnostic of the conservation status of exploited fish species. This simple and cost-effective tool can help prioritize research, management, and conservation on species with suspected loss of genetic diversity potentially eroding their adaptive potential to global change.

## Background

The exploitation of species’ wild populations can produce declines in population sizes, and even drive species to extinction (Hutchings & Reynolds 2004; Allendorf et al. 2008). Such exploitation has the potential to cause three types of genetic change: alteration of population structure and connectivity, selection induced genetic changes, and loss of genetic diversity (Allendorf et al., 2008; Gandra et al., 2021). As genetic diversity is the raw material for natural selection allowing species to adapt to new environmental conditions, its loss will decrease species adaptive potentials (Spielman et al., 2004; Conover et al., 2006; Allendorf et al., 2008; Hare et al., 2011; Pinsky & Palumbi et al., 2014; Bernatchez et al., 2017; Gandra et al., 2021). Hence, exploited species with eroded genetic diversity can be threatened by the loss of their adaptive potential to respond to changes in their environments.

Fish species have been exploited for several decades decreasing their population abundances and changing their age composition (Hutchings & Reynolds 2004; FAO 2020). However, the evidence of loss of genetic diversity by this exploitation has been rather contradictory with some studies showing decreased genetic diversity in exploited species (e.g. Hauser et al., 2002; Pinsky & Palumbi, 2014) but others did not (e.g. Pinsky et al., 2021). Pinsky & Palumbi (2014) addressed the question of the loss of genetic diversity in exploited fish species by a meta-analysis comparing genetic diversity estimates based on microsatellite genetic markers from exploited and non-exploited species. They found clear evidence of allelic richness loss in highly exploited fish populations. Moreover, Gandra et al. (2021) found a significant correlation between fish commercial importance and genetic structure, suggesting that commercially exploited species exhibit lower levels of genetic differentiation among populations. Nonetheless, genetics is rarely taken into account in fisheries management with the potential outcome of a hidden loss of adaptive potential to environmental changes in exploited fish species (Hauser & Carvalho, 2008, Allendorf et al., 2014; Pinsky & Palumbi, 2014; Ovenden et al., 2015; Casey et al., 2016; Bernatchez et al., 2017; Bryndum-Buchholz et al., 2021; Ovando et al., 2021; Gandra et al., 2021). Implementation of genetic evaluations for fish stocks in fisheries has been challenging. Reasons for this are that studies with more traditional microsatellite markers can be laborious and while new sequencing technologies are powerful to address these subjects (Hauser & Seeb, 2008; Bernatchez et al., 2017), specific training is required and high-throughput sequencing is costly, hence limiting the number of species to be studied.

Mitochondrial genetic markers have been used for phylogenetic and population genetics studies for many years. And more recently the use of Cytochrome Oxidase subunit I (COI) mitochondrial gene has widely increased for species identification (Ratnasingham & Hebert, 2007). Although the extensive use of this marker has raised some criticism because mitochondrial genome evolution has different characteristics than the nuclear genome such as haploidy and uniparental inheritance, the use of mitochondrial genetic markers gives very valuable information for under-studied species (Rubinoff & Holland, 2005). Whilst genetic drift and the mutation rate per generation are the two major forces driving both nuclear and mitochondrial genome evolution, the contributing forces of these two processes are different between the two genomes, with genetic drift being more important in nuclear and mutation rate in mitochondrial genomes (Lynch et al., 2006). However, if the differences and limitations of mitochondrial molecular markers are taken into account, their use can provide preliminary insights into population changes. For example, declines in population sizes are expected to decrease genetic diversity in both, mitochondrial and nuclear genomes as exhibited in threatened mammal species (e. g. Casas-Marce et al., 2017; van der Valk et al., 2018). Therefore, despite uncertainties on how much mitochondrial genetic variation can inform on the overall evolutionary history of the species, it can inform on demographic changes, and especially on declines in population sizes. Petit-Marty et al. (2021) demonstrated that the level of genetic diversity in the COI mitochondrial gene can be used as a proxy of the species′ conservation status. By using a curated dataset of COI genetic diversity estimates for more than 4000 animal species, significant differences were revealed between threatened and non-threatened animal species assessed by the International Union of Conservation of Nature (IUCN). After accounting for many biases, differences in levels of genetic diversity in COI were associated with declines of population census in threatened species, because species assessed as threatened in IUCN show evidence of global decline whereas non-threatened species do not. Differences were especially significant for 1426 worldwide distributed fish species (Petit-Marty et al., 2021). Additionally, a positive correlation between mitochondrial and nuclear genetic diversity has been found for fish species (Piganeau & Eyre-Walker, 2009). Thus, genetic diversity loss in mitochondrial genomes due to population declines correlates with the loss of genetic diversity in the nuclear genome and can therefore, inform on the conservation status of fish species populations.

Fisheries management, including fishing restrictions, varies widely across the globe (FAO, 2020; Ovando et al., 2021; Bryndum-Buchholz et al., 2021) as does the effect of fishing pressures, leading to the need for local assessments of fish populations. Data of stocks abundance and age composition for economically important species have been documented for decades (FAO 2020; Ovando et al., 2021), yet fisheries statistics are related to short-term reactions to exploitation. Genetic analyses, on the contrary, can inform on the long-term processes which are related to species adaptive potential to respond to changes in environmental conditions (Hauser & Carvalho, 2008; Pinsky & Palumbi, 2014; Bernatchez et al., 2017; Gandra et al., 2021). Ideally, the study of genetic diversity loss in wild populations would require samples from before and after species exploitation, however, such data is mostly not available for commercial fish species. Therefore, an alternative approach is to use comparative datasets of genetic diversity in related species. In this sense, the COI mitochondrial gene has the advantage to be widely studied as a barcode sequence in many species for their identification (Ratnasingham & Hebert, 2007). Therefore, hundreds of thousands of fish COI sequences are available in public databases (e.g. GenBank and Barcode of Life Data (BOLD) System), which can be used as comparative frameworks to assess the loss of genetic diversity under exploitation scenarios. Moreover, given its wide use as a genetic barcode, this molecular marker has the potential to be easily implemented in fisheries evaluations as getting sequences of many species does not require advanced training or huge budgets. Here we evaluated the levels of genetic diversity in COI mitochondrial gene for fish species with different degrees of commercial importance. These fish species were collected from the East China Sea where there is evidence of an elevated exploitation (Chang, Shiao & Gong 2012; Liang & Pauly, 2017; Zhang et al., 2018; Sumaila, 2019; Teh et al., 2020; Zhang et al., 2020). The genetic diversity levels were contrasted against large datasets of fish species. The comparisons were calibrated by adding two farmed species with expected low genetic diversity, and four species with long-term fishing management and extensive knowledge on their population dynamics. The results indicate that this molecular marker could be used for an approximation to the conservation status of fisheries and a valuable guide to prioritizing further case-study research.

## Methods

### Sampling and species identification

Fixed nets were settled in 6 different locations within the Sansha Bay in Nindge, Fujian Province of China (East China Sea, Latitude: 119°43′-119° 49′; Longitude: 26°34′-26°41) once a month (full or new moon phase) from April to September of 2019. Fin clips were stored in alcohol 95% for the collected fish species. We took the ten fish species with accumulated sample sizes larger than 30 individuals as representatives of the most abundant fish in Sansha Bay (*Collichthys lucidus*, *Crenimugil crenilabis*, *Cynoglossus oligolepis*, *Harpadon nehereus*, *Larimichthys crocea*, *Siganus fuscescens*, *Stephanolepis cirrhifer*, *Takifugu oblongus*, *Thryssa vitrirostris*, and *Trypauchen vagina*). One of the species, *Larimichthys crocea*, is not representative of wild populations but farmed resulting from long-term (since 2002), restocking activities in Sansha Bay after the natural populations were depleted in the 1980s (Liu & Sadovy de Mitcheson 2008, Liu et al. 2020, Yuan et al. 2021), and it was taken as a negative control (i.e. expected low genetic diversity) in the evaluation of the other fish species. We also processed samples from another farmed species collected in Hong Kong waters, the hybrid Sabah grouper (*Epinephelus fuscoguttatus* females *x E. lanceolatus* males), which was also used as a negative control in the evaluation analysis. Additionally, we processed samples of four highly commercial fish species (*Lophius budegassa*, *Merluccius merluccius*, *Mullus barbatus* and *Mullus surmuletus*) from Balearic Islands (western Mediterranean, Latitude: 38°50′-40°50′; Longitude: 2°00′-4°50′) which present long-term managed fisheries and fishing restrictions. These species were collected during the MEDITS bottom trawl surveys in 2019, which are carried out annually during spring and early summer trough the Balearic Islands (for sampling protocol see Spedicato et al. 2019).

DNA was extracted from fin clips for 24 samples of the ten representative species from Fujian, 20 samples of the hybrid grouper from Hong Kong and among 19-45 samples from the four species of Balearic Islands using the DNAeasy blood and tissue Qiagen kit. COI mitochondrial gene was amplified by PCR following the protocol of Ivanova et al. (2007), with a final volume of 15 ul and 4 ul of DNA (1ng/ul) using the primers cocktail COI-3. PCR products were purified with the Qiagen PCR purifying kit and sequences were obtained by a Sanger sequencer from the Centre of PanorOmic Sciences of the University of Hong Kong using the same primers as in the PCR and 6-12 ng of purified PCR products. Sequences with average phred scores > 50 and samples sizes ≥ 19 were successfully achieved for nine out ten of the more abundant species collected in Fujian (i.e. all but *C. oligolepis*), the hybrid grouper, and the four species from Balearic Islands. All species were taxonomically identified by an expert and species identity by COI sequences was confirmed in BOLD system (http://www.boldsystems.org, Ratnasingham & Herbert 2007). The taxonomic identity of the species was the same for the visual and molecular identification for all but one, *S. fuscescens* for which the identification at species level was not possible in BOLD system and therefore it was not included in further analyses.

Sequences were aligned using Muscle software (Edgar, 2004) and genetic diversity estimates (the average number of nucleotide differences per site between two sequences, π, Nei, 1987), Tajima’s D (Tajima, 1989) and isolation index, F_ST_, were obtained using DNAsp software (Rozas et al., 2017). We used π (Nei, 1987) genetic diversity estimates (COI-π) instead of ⍰ (Watterson, 1975) because the former is based in frequency and therefore, a better indicator of demographic changes (Tajima, 1989). Two different clades have been recognized by DNA barcoding in *Trypauchen vagina* (Thu et al., 2019), accordingly, the sequence identification in BOLD systems shows the presence of the two clades among our sequences. The two clades are genetically different (F_ST_ =0.97, p_Fst_=0.018), and therefore, we only present the results for the major clade found in our dataset of *T. vagina* (N=15). Variation in estimates of genetic diversity based on samples sizes larger than 15 is lower than < 5e^-4^ in COI for species with COI-π < 0.01, (Petit-Marty et al., 2021), and thus, our estimates can be considered representative of the population. All sequences are deposited in NCBI nucleotide database with accession numbers: *Collichthys lucidus*: <u>OL673515-OL673535</u>; *Crenimugil crenilabis*: <u>OL673809-OL673829</u>; *Harpadon nehereus*: <u>OL673927-OL673947</u>; *Larimichthys crocecr*: <u>OL674152-OL674171</u>; *Stephanolepis cirrhifer*: <u>OL684530-OL684553</u>; *Takifugu oblongus*: <u>OL684869-OL684890</u>; *Thryssa vitrirostris*: <u>OL679101-OL679121</u>; *Trypauchen vagina*: <u>OL684325-OL684339</u>; hybrid Sabah grouper: <u>OL674054-OL674073</u>; *Lophius budegassa*: <u>OL674085-OL674103</u>; *Merluccius merluccius*: <u>OL684344-OL684384</u>; *Mullus barbatus*: <u>OL6848910L684915</u>; and *M. surmuletus*: <u>OL674197-OL674229</u>.

### Data analyses

To investigate the confidence intervals (CIs) of COI genetic diversity values (COI-π) we used the dataset of fish species obtained by Petit-Marty et al. (2021) as a comparative framework (N=1426). We also estimated more conservative 95% CIs filtering the worldwide dataset for species distribution (East Asia and Mediterranean) and habitats (Marine and Estuaries species for East-Asia, and Marine for Mediterranean), and eliminating COI-π estimates based on sample sizes of N<15. Because the worldwide dataset is representative of the whole species, and genetic differentiation among populations within species could give COI-π not representative of within-population genetic variation, we also eliminated the species with COI-π estimates higher than the upper boundary of the mean 95% CI of the worldwide dataset for a more conservative final datasets. The list of species and their estimated genetic diversity values used for the comparative frameworks of East-Asia and Mediterranean is presented in **Suppl. Table S1.** 95% Confidence Intervals (CI) of the median and mean values of genetic diversity in COI were obtained by the boot package (Hesterberg, 2011) in R by bootstrapping 10,000 datasets of the mean and median values of π. Overall trait data on the species, such as commercial importance, body size, geographic distribution, and generation times for the East Asia species were retrieved from Fishbase (https://www.fishbase.se; Froese & Pauly, 2021), IUCN species assessments (https://www.iucnredlist.org), and references of studies from individual species and fisheries assessments. Population trends and habitat information for Balearic Islands species was obtained from local assessments (Guijarro et al., 2012; 2019a; 2019b), with the exception of *L. budegassa*, for which the Central Mediterranean data was used, the closest area where this stock has been assessed (Fiorentino et al., 2012).

Species were classified for their commercial importance as highly, minor, and not commercial. We considered highly commercial species those under fishing pressures in their whole distribution and traded in local, national, and international markets. Minor commercial are species that are traded only in local and/or national markets and therefore, their levels of exploitation can vary across its distribution. Finally, not commercial are species not suitable for human consumption.

### Simulation analyses

We used msms program to generate samples under a Wright-Fisher neutral model of genetic variation (Gutenkunst et al., 2009) simulating declines of population sizes. We evaluated a simple model that does not account for the complex dynamics of fish populations, but enhances the expectations of levels of genetic diversity under the effects of continued harvesting. In the simulations, the populations experienced a decline in population sizes between 1e^6^ and 1 generations ago, with no recombination, no migration, and no further population expansion after declines. We used a value of starting genetic diversity of ⍰=2.5 (~Neμ; Watterson, 1975) which is equivalent to the median value observed for East Asia and worldwide fish species datasets **(Table 1)**. We tested different effective population sizes (1e^6^, 1e^5^, and 1e^4^) and evaluated the average values of the average pairwise nucleotide differences among sequences of 650 base pairs for 1000 random samples of 20 individuals and different degrees of declines in population sizes (10%, 30%, 50%, and 90%).

**Table 1.** 95% Confidence Intervals (CI) of median and mean values of COI genetic diversity (COI-π) for datasets of fish species. East-Asian and Mediterranean datasets are filtered subsets of worldwide dataset (see Methods for details).

| Dataset | No. of species | 95% tCI Median | 95% CI Mean |
| --- | --- | --- | --- |
| World-wide | 1426 | 0.0035-0.0040 | 0.0101-0.0118 |
| East Asia | 118 | 0.0035-0.0049 | 0.0041-0.0051 |
| Mediterranean | 58 | 0.0028-0.0042 | 0.0034-0.0047 |

## Results

The first step of our analysis was setting up the comparative frameworks based on worldwide datasets of genetic diversity in COI estimates for fish species. The 95% Confidence Intervals (95%CI) of mean and median values for a worldwide dataset of fish species (N= 1426) did not overlap between threatened (i.e. assessed as VU, EN, and CR in IUCN, N=112) and non-threatened fish species (i.e. assessed as LC at IUCN, N=1260) as previously showed in Petit-Marty et al. 2021. However, mean values are biased to high values owing to the not normal distribution of COI-π **(Suppl. Figure 1).** Thus, for species evaluations, we used the more conservative 95% CI low boundary of median values calculated for worldwide fish species that coincide with the East Asia dataset **(Table 1)**. 95% CI of median values for Mediterranean species show slightly lower values, likely representing historical events such as the last glaciations event **(Table 1)**.

The evaluation of the species from Fujian (East China Sea) shows that four out of seven species (the two highly commercial: *Harpadon nehereus*, and *Stephanolepis cirrhifer*; and two minor commercial: *Crenimugil crenilabis*, and *Trypauchen vagina*,) exhibit COI-π below those expected for 95%CI of the median for worldwide and East Asian species **(Figure 1, Table 2)**, and within the expectations for world-wide threatened species **(Suppl. Figure 1)**. The non-commercial fish *Takifugu oblongus* was the only species presenting COI-π levels higher than expected by the upper boundary of the 95% CI of the median for East-Asian species **(Figure 1)**. Accordingly to our hypothesis, loss of genetic diversity in COI could be indicative of population declines eroding the adaptive potential of the species populations. Therefore we classified the four species from Fujian with detected low values of COI-π as First Priority species for further research on their populations’ conservation status **(Figure 1)**.

**Figure 1.**
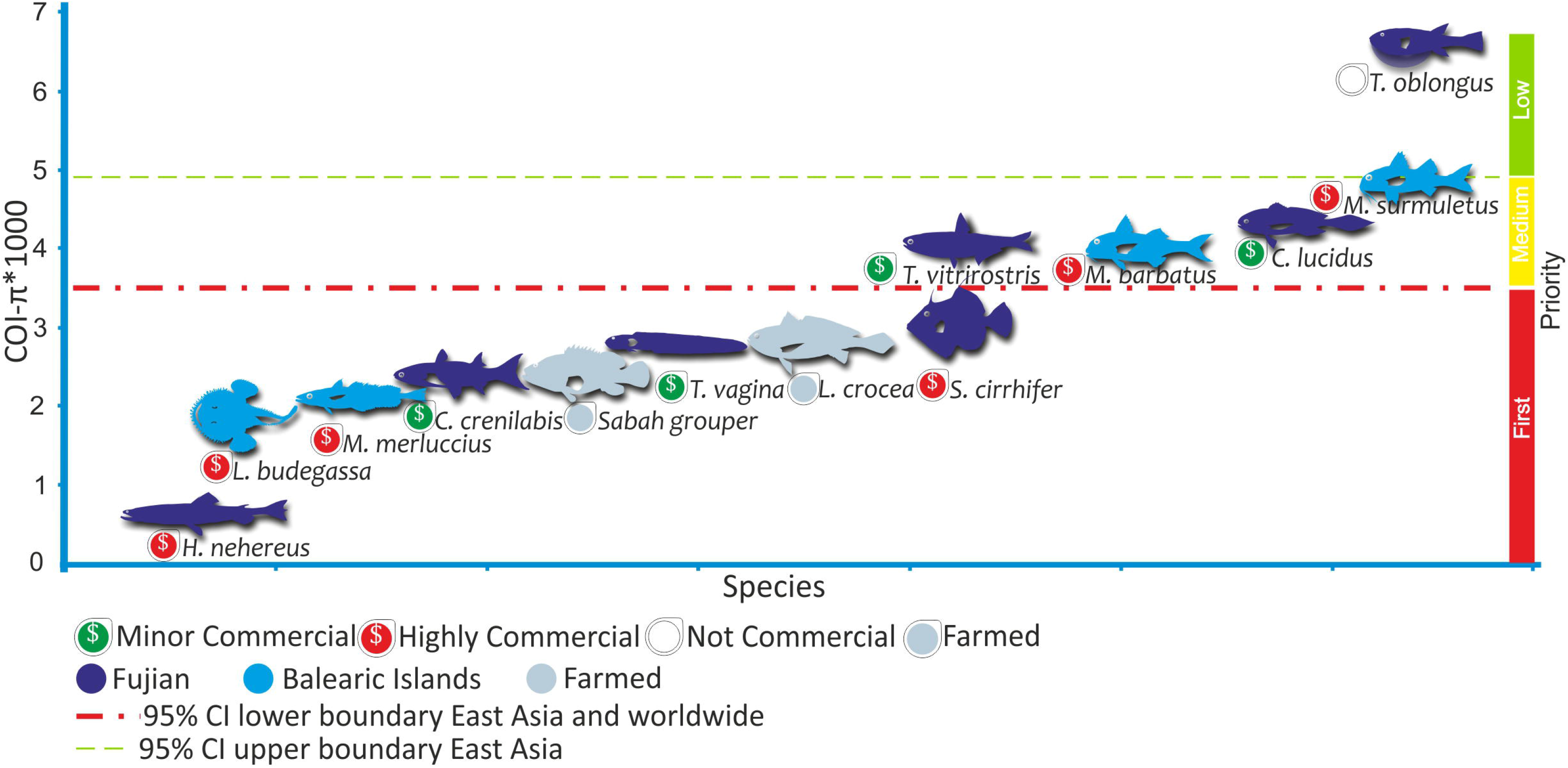
Levels of genetic diversity in COI mitochondrial gene (COI-π) of fish species. Species were ordered by their levels of COI-π and these were multiplied by 1,000 for graphical purposes. The red line shows the lower boundary of the 95%CI of the distribution of median genetic diversities values obtained by bootstrapping worldwide and East Asian fish datasets. The green line shows the upper boundary of the 95%CI of the median distribution of genetic diversity in East Asian fish. Fish from Fujian are the evaluated species in this study. Fish from the Balearic Islands are long-term managed species with known population dynamics. Farmed fish are expected to have low levels of genetic diversity and were used to calibrate the comparisons. Highly commercial species are under fishing pressures in their whole distribution while fishing pressures are variable for minor commercial species. Illustration credits: Gaston Petit.

**Table 2.** Sampled fish species, their characteristics and genetic diversity estimates in COI mitochondrial gene **(COI-π).** Max.size indicates maximum size registered in cm, gen.times is generation time in years, IUCN: IUCN assessments, Pop trends: populations trends, D: Tajima’s D statistics; N: analysed sample size. nd: no data

| Species | Commercial importance | IUCN | Pop. trends | Threats | Movements/Habitat | Max.size (cm) /gen.times (year) | COI-π | D | N |
| --- | --- | --- | --- | --- | --- | --- | --- | --- | --- |
| **Evaluated species** |  |  |  |  |  |  |  |  |  |
| *Harpadon nehereus* | high | NT | decreasing | Overfishing | Oceanodromous/<br>Marine-Neritic, subtidal, estuaries | 40/3y | 0.0002 | -1.16 | 21 |
| *Stephanolepis cirrhifer* | high | LC | stable | Overfishing | Oceanodromous/<br>Marine-Neritic, subtidal | 30/nd | 0.0027 | -2.13* | 24 |
| *Crenimugil crenilabis* | minor | LC | unknown | Overfishing/<br>by-catch | Non-migratory/<br>Marine-Neritic, coral reef, estuaries;<br>wetland (inland) | 60/nd | 0.0020 | -1.99* | 21 |
| *Thryssa vitrirostris* | minor | LC | unknown | Overfishing/<br>by-catch | nd/<br>Marine-Neritic, pelagic, estuaries | 18/1y | 0.0035 | 1.11 | 21 |
| *Trypauchen vagina* | minor | LC | unknown | Overfishing/<br>by-catch | Amphidromous/<br>Marine-Neritic, , coral reef, estuaries;<br>Marine-intertidal, mud, mangroves;<br>wetland (inland) | 18/1y | 0.0024 | -1.21 | 15 |
| *Collichthys lucidus* | minor | LC | unknown | Overfishing/<br>by-catch | Oceanodromous/<br>Marine-Neritic, subtidal, estuaries | 17/1y | 0.0040 | -2.07* | 21 |
| *Takifugu oblongus* | not | LC | decreasing | Unknown | nd/<br>Marine-Neritic, subtidal, estuaries<br>Marine-intertidal, mangroves | 40/nd | 0.0062 | -1.75 | 23 |
| **Control farmed species** |  |  |  |  |  |  |  |  |  |
| *Larimichthys crocea* | Mostly farmed - | - | - | - | Oceanodromous<br>Marine-Neritic, subtidal, estuaries | 80/15y | 0.0024 | -2.18* | 20 |
| *Sabah grouper*<br>(*Epinephelus fuscoguttatus* x<br>*E. lanceolatus*) | Farmed | - | - | - | Oceanodromous<sup>a</sup><br>Marine-Neritic, coral reef, seagrass,<br>estuaries<sup>a</sup> | >200/25.5y<sup>a</sup> | 0.0020 | -2.14* | 20 |
| **Control species from long-term managed fisheries in Balearic Islands** |  |  |  |  |  |  |  |  |  |
| *Merluccius merluccius* | high | VU | Oscillating | Overfishing | Low migrant | 82 /9y | 0.0018 | -0.58 | 41 |
|  |  |  |  |  | Demersal (deep shelf and upper slope) |  |  |  |  |
| *Lophius budegassa* | high | LC | Oscillating | Overfishing | Low migrant<br>Demersal (deep shelf and slope) | 90 /10y | 0.0016 | -2.20* | 19 |
| *Mullus surmuletus* | high | LC | Decreasing | Overfishing | Oceanodromous<br>Demersal (Shallow and deep continental shelf) | 39 /4.1y | 0.0047 | -1.73 | 34 |
| *Mullus barbatus* | high | LC | Decreasing | Overfishing | Low migrant<br>Demersal (Shallow and deep continental shelf) | 30 /3.6y | 0.0035 | -0.85 | 25 |

\*p<0.05; <sup>a</sup> based on *E. fuscoguttatus*

We contrasted the levels of COI-π of these four First Priority species to those of two farmed species *Larimichthys crocea* and the hybrid Sabah grouper finding that the levels of COI-π in the farmed is similar or higher than the First Priority species **(Figure 1, Table 2).** Both farmed species very likely experienced a decrease in their levels of genetic diversity by the founder effect when starting the farms (Wang et al., 2012). Moreover, wild populations of *L. crocea* were assessed as critically endangered (Liu et al. 2020), while *E. fuscoguttatus* (female of the hybrid, Rhodes et al., 2018) had been assessed as vulnerable, and therefore both species presented evidence of decline in populations sizes >30% (i.e. the cut-off used in IUCN to classify a species as threatened) before starting farming. Accordingly, the significant negative value of Tajima’s D **(Table 2)** in these two farmed species shows evidence of a recent bottleneck (Tajima, 1989). Similarly, four other species show significant negative values of Tajima’s D (*S. cirrhifer*, *C. crenilabis*, *C. lucidus*; and *L. budegassa* **Table 2)**. However, lack of significance in Tajima’s D does not rule out ongoing populations’ declines producing very low genetic diversity values in both estimators used to calculate D, π and ⍰ (Tajima, 1989). Thus, when comparing COI-π levels of the farmed species against the worldwide COI-π 95% CI they show reduced genetic diversity as expected by founder events when the farms were started, indicating that decline in population sizes can explain also the low COI-π in First Priority species.

Most of the evaluated species from Fujian have short body sizes (i.e. ≤40 cm of maximum length, **Table 2)**, which could be taken as indicative of short generation times (≤ 3 years, for species with known generation times, **Table 2).** Therefore, in terms of generation times, the evaluated species can be compared to the two species with short generation times and managed populations from Balearic Islands (*Mullus barbatus*, and *M. surmuletus*; **Table 2).** We found that the two species of *Mullus* from Balearic Islands presented levels COI-π within the expected for the worldwide dataset **(Figure 1)**, with *M. sumurletus* showing COI-π levels even above of the upper 95%CI boundary of the Mediterranean species, despite a tendency to declining populations found for both species (Guijarro et al., 2012; 2019a). Thus, for species with short generation times, the long-term management and the fishing restrictions in the Balearic Islands seem to help maintain average levels of COI-π, while overfishing in East China Sea could increase the loss of genetic diversity in small species.

On the contrary, the other two long-term managed species from Balearic Islands, *M. merluccius* and *L. budegassa*, are larger with longer generation times (~10 years, **Table 2)** and show levels of COI-π below the low boundary of 95%CI for Mediterranean and worldwide species **(Figure 1, Table 1)**. Therefore, the comparison between small and large species from Balearic Islands (i.e. *M. barbatus*, *and M. surmuletus* vs *M. merluccius*, and *L. budegassa*, **Table 2)** suggest that differences in generation times may account for the observed differences in COI-π in long-term managed species.

Finally, we used coalescent simulations to find out what are the expectations in decreases of genetic diversity by population declines under a neutral model of evolution. The results of such simulations show that under a simple neutral model of molecular evolution and population’s declines, with no recombination, no migration and no further expansion, decreases in the average levels of genetic diversity are proportional to the decline in population census **(Figure 2).** The probability to find values of genetic diversity ≥ 2.5 (i.e. starting value of simulations) among 1000 generated neutral samples was p<0.005, ~0.10, ~0.20, and ~0.40 for declines in populations of 90%, 50%, 30%, and 10%, respectively. The simulations also indicate that the observed levels of genetic diversity in the species classified here as First Priority **(Figure 1)** could indicate declines in population sizes of ~40% or more, but *H. nehereus*, could have experienced stronger declines of nearly 90% of its population.

**Figure 2.**
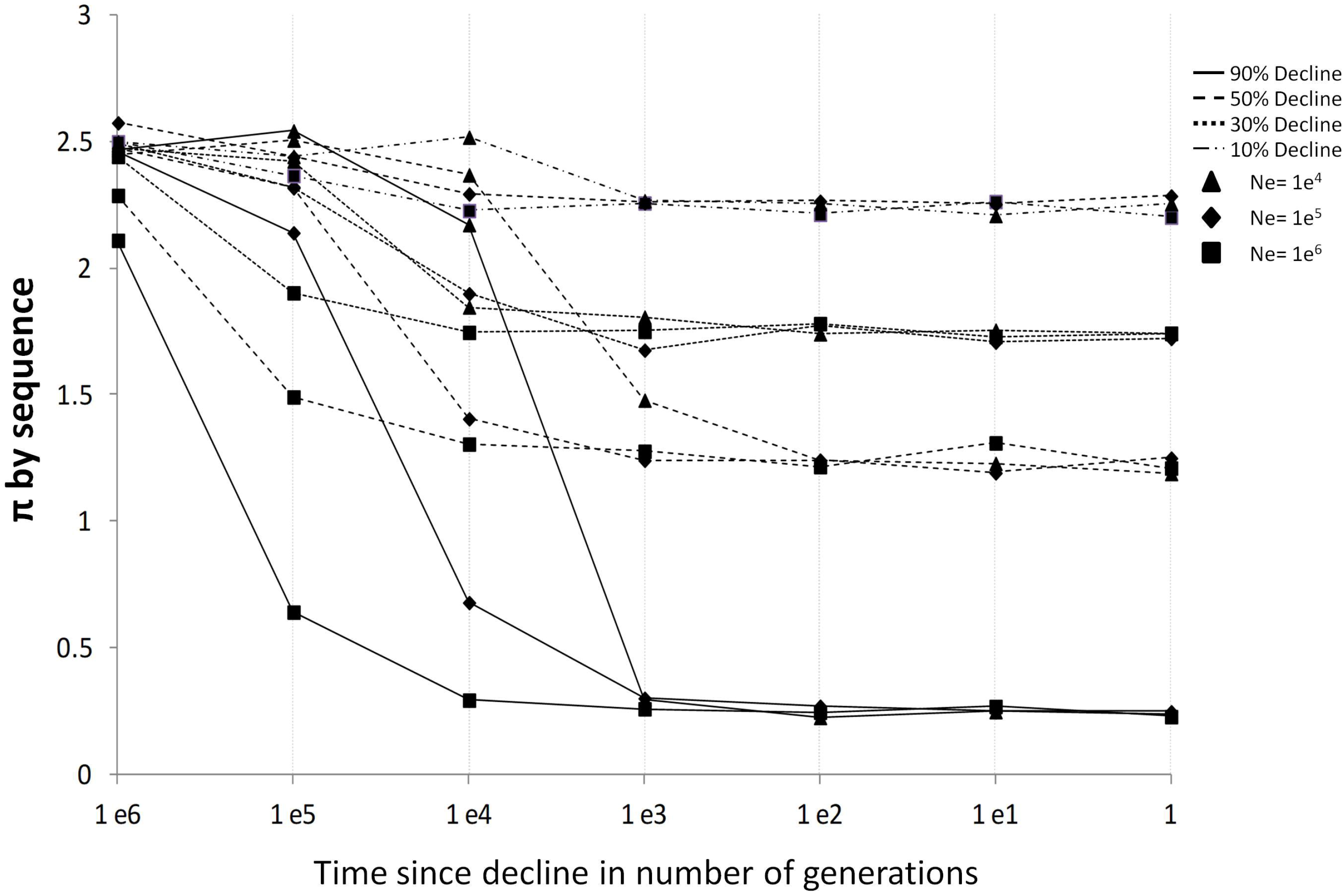
Simulations of the effect of populations declines of the 10%, 30%, 50%, and 90% of its original size on genetic diversity. The starting value of ⍰ was 2.5, which is similar to the median value of worldwide and East Asia fish datasets **(Table 1).** Note that variations in Ne for a fixed ⍰~=Neμ indicate variations in mutation rates. The number of generations when the decline happened is scaled to Ne, and therefore no effects of decline are detected when this happened ~Ne generations ago.

## Discussion

The results of contrasting the levels of COI genetic diversity (COI-π) of the fish species collected in Fujian (East China Sea) against expected average levels for worldwide and East Asian species suggest loss of genetic diversity owing to fishing pressures. The only species with levels of genetic diversity above the expected average for East-Asia and worldwide species is *Takifugu oblongus* which is a poisonous species and not consumed in East China (Shao et al., 2014). On the contrary, the two highly commercial species *H. nehereus* and *S. cirrhifer*, and two minor commercial species, *T. vagina* and *C. crenilabis* show reduced levels of genetic diversity below the expected average for East-Asia and worldwide species. These results are in agreement with evidences of threats found in assessments of these species at IUCN. Evidence of over-exploitation has been found for the two highly commercial species, *H. nehereus*, and *S. cirrhifer* (Matsuura et al., 2019; Russell et al., 2019), while artisanal fishing and by-catch are major threats for all minor commercial species evaluated in this study with the only exception of *C. crenilabis* which is under-studied (Hoese & Sparks, 2017; Dinh, 2018; Larson, 2019, Hata 2020; Nguyen Van et al., 2020, Zhang et al. 2020). Thus, the first result emerging from our comparative analyses indicates that both high and minor commercial species from Fujian (East China Sea) have potentially experienced decreases in their levels of genetic diversity by fishing pressures.

Beyond declines in population census by exploitation, the levels of genetic diversity are also affected by biological, ecological and evolutionary history characteristics of the species. Genetic diversity is defined as the product of effective population size (Ne) by mutation rate per generation (μ). Mutation rates per generation can be considered similar when using the same molecular marker across related species and are unlikely to explain the observed differences in COI-π (Petit-Marty et al., 2021). Nonetheless, the differences in long-term effective population size (i.e. the effective population size of the species in equilibrium before demographic changes, Ne) and generation times among species, may account for the observed differences, and we discuss these alternative explanations to declining population size.

A first alternative explanation to the decline of population size for the observed levels of COI-π of the evaluated fishes is that Ne would be small in the evaluated species. Although COI-π has been shown to be a good indicator of population declines (Petit-Marty et al., 2021), there are uncertainties on how informative mitochondrial variation is on the long-term effective population size (Ne; Lynch et al., 2006). However, Piganeau & Eyre-Walker (2009) found evidence of a positive correlation between mitochondrial and nuclear genetic diversities estimates in fish. Moreover, body size and generation times are likely positively correlated, while body size correlates negatively with Ne (Lynch & Connery, 2003; Pinsky & Palumbi, 2014). Thus, if COI-π is correlated with species Ne, then it could be expected that species with large body sizes and generation times and, in turn, expected low Ne, have low variation in COI-π. Most species evaluated here, have small sizes (≤ 40 cm maximum length) and their generation times are likely to be short (<5 years). However, the two farmed species which have large body sizes and long generation times show levels of COI-π similar or even higher than the species classified as First Priority here. Moreover, when comparing the evaluated species to the two species with short body sizes and generation times from Balearic Islands, *M. barbatus* and *M. surmuletus*, we found π-COI levels of >40% higher in Balearic Islands species than in the First Priority, and farmed species. Hence, these comparisons suggest that levels of COI-π found in the First Priority species is more likely a result of declines of population census than differences in Ne.

A second alternative explanation for the observed differences in genetic diversity could be due to differences in generation times. The neutral theory of molecular evolution explains that the observed levels of genetic diversity account for events (demographic or selective) that happened between 1 and ~Ne generations ago (4Ne for nuclear markers, Lynch & Connery 2003). This implies that for species with long generation times the recovery time of genetic diversity in the population will be at least several hundreds or thousands of years, assuming no population expansion after a decline (e.g. if fishing pressures are constant across time) and restricted rates of migration. Moreover, it may imply that for species with long generation times, as is the case for two large species from the Balearic Islands *L. budegassa* and *M. merluccius* (~10 years), if the effective population size is >~1e^3^, the levels of genetic diversity in these species may account for past evolutionary events as the last glacial period (Pinsky & Palumbi, 2014). In fact, *L. budegassa* shows a significant negative Tajima’s D value suggesting a past bottleneck, as regional assessment for this species does not indicate declines in the population census for the last three decades (Papakonstantinou et al., 2011; Fiorentino et al., 2012). On the contrary, the populations of *M. merluccius* from the Mediterranean Sea were regionally assessed as vulnerable by evidence of declines in population census (Di Natale et al., 2011; Guijarro et al., 2019b) and exhibit fine-scale population structure within the Mediterranean populations (Milano et al., 2014) likely affecting local levels of genetic diversity. Both species, *L. budegassa* and *M. merluccius*, are of high commercial importance and have been under fisheries management measures for at least three decades, with no evidence of population increase or recovery (Di Natale et al., 2011; Papakonstantinou et al., 2011; Fiorentino et al., 2012; Guijarro et al., 2019b). Moreover, *L. budegassa* shows slightly lower values of genetic diversity (0.0016) than *M. merluccius* (0.0018), while the census of their local populations shows inverse trends. In the Balearic Islands, *M. merluccius* has high abundance values (26325 ind. km^2^; Sion et al., 2019), while *L. budegassa* is less abundant (183 ind. km^2^; Barcala et al., 2019). Hence, while declines in population sizes and population structure could explain low genetic diversity in *M. merluccius*, the reduced level of genetic diversity in *L. budegassa* could be explained by smaller long-term effective population size (Ne) and past bottlenecks. Thus, the levels of COI-π for *L. budegassa* and *M. merluccius* are not reflecting the effect of fishing pressures alone, but more likely the combination of continuous harvesting with past population bottlenecks, the effect of long-term fisheries management changing population structure and connectivity, and fishing induced selection (Allendorf et al., 2008; Gandra et al., 2021). Therefore, comprehensive population genomics studies might be needed in order to disentangle the effect of harvesting by itself on the loss of genetic diversity for large species.

Finally, species characteristics such as large variability in reproductive success, unequal sex-proportion, connectivity among populations, age and stage-structured populations, and habitat preferences have been found affecting the levels of genetic diversity in fish (Hauser & Carvalho, 2008; Hare et al., 2011; Pinsky & Palumbi, 2014; Eldon et al., 2016; Martinez et al., 2018; Gandra et al., 2021). The power in this study is not sufficient to evaluate all biological and ecological traits affecting genetic diversity, but we can have a preliminary evaluation of differences in connectivity among populations and habitat preferences.

Migration is a likely source of genetic variation in fish populations from both, adults and pelagic larval stages, counteracting the effects of local declines (Hauser & Carvalho 2008; Pinsky & Palumbi, 2014; Martinez et al., 2018). All evaluated species are non-guarders with pelagic larvae, thus, recruitment of larvae from different populations could also increase local genetic diversity. Among the evaluated species, the First Priority *T. vagina*, is known to have elevated rates of self-recruitment (Dinh, 2018), suggesting that low rates of dispersion of pelagic larvae could increase the losses of genetic diversity for minor commercial species. Moreover, three evaluated species from Fujian are known to be oceanodromous (i.e. seasonally migratory fishes). These species have different commercial values (i.e. Minor: *C. lucidus vs* High: *H. nehereus* and *S. cirrhifer*) and we observed the highest levels of COI-π in the species with minor commercial importance. This comparison suggests that for the Medium Priority species *C. lucidus* with minor commercial importance and likely variability in fishing pressures across its distribution, the levels of genetic diversity could be maintained owing to migration. However, for the highly commercially important species (*H. nehereus* and *S. cirrhifer*), on the other hand, which are under fishing pressures throughout its distribution range, the levels of genetic variation are likely decreasing globally for the whole species. Moreover, low COI-π observed for the non-migratory species *C. crenilabis* (First Priority) also suggests that migration may account for the differences in COI-π levels in minor commercial species. Therefore, by taking into account migration rates of the species, it is suggested that for highly commercially important unmanaged species in Fujian the detected decrease in genetic diversity in COI seems to be directly linked to their global fishing pressures. In contrast, for minor commercial species, differences in migration rates may account for the differences in genetic diversity.

The two *Mullus* species from the Balearic Islands also differ in their migration rates, with the highest COI-π in the migratory species *M. surmuletus*. However, the differences in the levels of COI-π between the two *Mullus* species may also be related to different habitat preferences. *M. surmuletus* prefers narrow shelf areas with rocky substrates while *M. barbatus* is more abundant in areas where shelf becomes wider with muddy bottoms (Lombarte et al., 2000; Tserpes et al., 2019). Because rocky areas are not accessible by bottom trawling, a significant proportion of the *M. surmuletus* population is not directly affected by this threat. Thus, when considering habitat preferences, the level of fishing pressures is also reflected in the genetic diversity levels of both long-term managed *Mullus* species.

Overall, the results of this study indicate that the differences in the levels of genetic diversity in COI for unmanaged exploited species might be explained by differences in fishing pressures causing declines in populations, and this empirical data is also supported by simulations. Although our simulations are simplistic, these show that the expected decrease in average genetic diversity in absence of migration and population expansion is proportional to the decline in population census. We found more than 80% of the simulated samples showing a decrease in the levels of genetic diversity when declines in population census were ≥30% (i.e. cut off to be considered as threatened under IUCN criteria). The best example of expected genetic diversity decreases with a decline in populations is *H. nehereus*,the species with the strongest evidence of unmanaged exploitation across its whole distribution (Russell et al., 2019) among the studied species. We found that the levels of genetic diversity in COI for *H. nehereus* could be explained by a decline in population size of more than 90%. In fact, though local assessment in Fujian has not been performed for this species, the decline in global population census was already estimated to be larger than 30%, with detected local declines of nearly 80% in Indonesia (Russell et al., 2019). Therefore, the observed genetic diversity in COI found in the evaluated species mostly reflects the reported declines in population census in the IUCN assessments, indicating that the use of COI-π is a good proxy for the conservation status of fish populations.

As genetic diversity in mitochondrial genome is higher than in nuclear genes (Lynch et al., 2006) using COI-π gives strong statistical power to detect potential genetic diversity loss than surveying genetic variation in the nuclear genome. For example, estimates of nuclear genetic diversity in cod from samples of more than 100 years ago were ~0.0008 (Pinsky et al., 2021) which would give few chances to detect genetic diversity loss using samples sizes of 20. Given the low economic cost of obtaining COI sequences, tens or hundreds of species could be locally evaluated with the same sampling effort, giving a clear first insight into the conservation status of local fisheries.

## Conclusions

Our results indicate that the estimates of genetic diversity in the mitochondrial gene COI account for: **1)** differences in fishing pressures by commercial importance of the fish species, **2)** theoretical predictions of decreases in genetic diversity produced by population decline, and **3)** expectations from local assessments of fisheries. The observed levels of COI-π suggest loss of genetic diversity in all fish species with high commercial importance, except long-term managed small-sized species which exhibit average levels of COI-π. Whilst the connectivity among populations could affect levels of genetic diversity for local populations of minor commercial species exploited locally in East Asia. Accordingly, we conclude that COI genetic diversity levels can provide a much-needed simple diagnostic of the conservation status of fish species under exploitation.

## Acknowledgements

Funding for this study were provided by the Area of Ecology & Biodiversity, School of Biological Sciences (HKU) to NPM and CS, the HKU start-up fund to CS, and by Fujian Province Ocean and Fisheries Bureau of China to ML. The MEDITS surveys are co-funded by the European Union through the European Maritime and Fisheries Fund (EMFF), within the National Program of collection, management and use of data in the fisheries sector and support for scientific advice regarding the Common Fisheries Policy. SRA was supported by a postdoctoral contract co-funded by the Regional Government of the Balearic Islands and the European Social Fund.

## Data Availability Statement

The data that support the findings of this study are openly available at NCBI nucleotide database at https://www.ncbi.nlm.nih.gov/nuccore, with reference numbers: *Collichthys lucidus*: <u>OL673515-OL673535</u>; *Crenimugil crenilabis*: <u>OL673809-OL673829</u>; *Harpadon nehereus*: <u>OL673927-OL673947</u>; *Larimichthys crocea*: <u>OL674152-OL674171</u>; *Stephanolepis cirrhifer*: <u>OL684530-OL684553</u>; *Takifugu oblongus*: <u>OL684869-OL684890</u>; *Thryssa vitrirostris*: <u>OL679101-OL679121</u>; *Trypauchen vagina*: <u>OL684325-OL684339</u>; sabah grouper: <u>OL674054-OL674073</u>; *Mullus barbatus*: <u>OL684891-OL684915</u>; *Merluccius merluccius*: <u>OL684344-OL684384</u>; *Mullus surmuletus*: <u>OL674197-OL674229</u>; and *Lophius budegassa*: <u>OL674085-OL674103</u>.

