## Supplementary for "Declining population sizes and loss of genetic diversity in commercial fishes: a simple method for a first diagnostic"

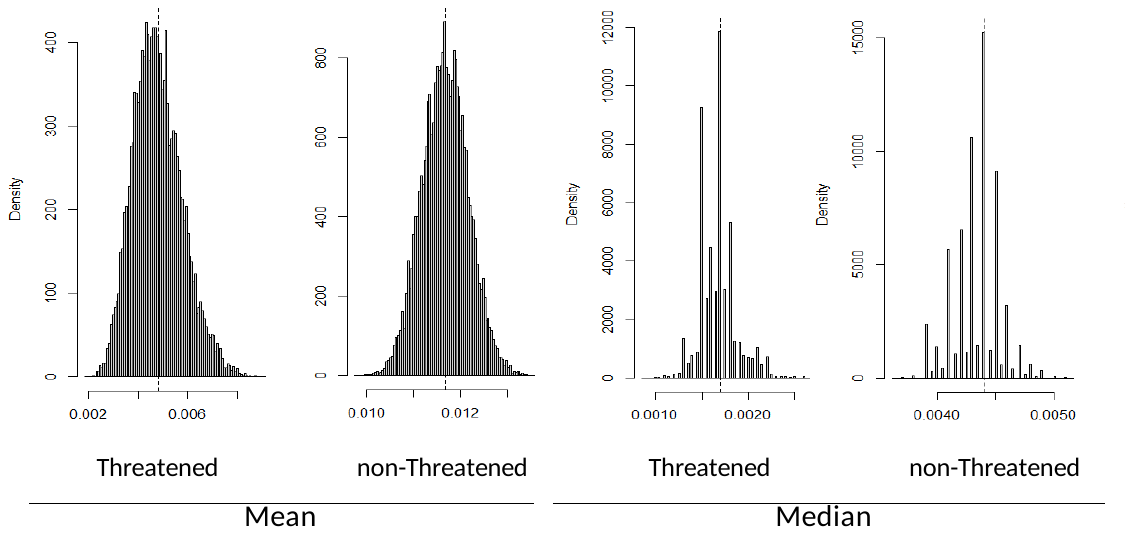

**Supplementary Figure S1.** Distribution of 10,000 bootstrapped COI genetic diversity values (COI-π) of the groups of threatened fish species (N=112, VU, EN and CR species in IUCN) and non-threatened species (N=1260, LC species in IUCN).

**Supplementary Table S1**. Comparative framework: Species datasets from East-Asia (N = 118) and Mediterranean (N = 46) with the sample numbers and estimated genetic diversity for the mitochondrial COI gene, obtained from Petit-Marty et al. 2021 and filtered according to geographical distribution and habitat as described in IUCN assessments.

| **Dataset** | **species** | **Nº individuals** | **π-COI** |
| --- | --- | --- | --- |
| East-Asia | Abudefduf vaigiensis | 239 | 0.0085 |
| East-Asia | Coryphaena hippurus | 190 | 0.0017 |
| East-Asia | Megalaspis cordyla | 119 | 0.0073 |
| East-Asia | Thunnus albacares | 118 | 0.0014 |
| East-Asia | Terapon jarbua | 98 | 0.0039 |
| East-Asia | Cephalopholis urodeta | 89 | 0.0033 |
| East-Asia | Cephalopholis argus | 81 | 0.0058 |
| East-Asia | Scomber japonicus | 77 | 0.0083 |
| East-Asia | Chaetodon collare | 59 | 0.0014 |
| East-Asia | Makaira nigricans | 59 | 0.0041 |
| East-Asia | Alepes kleinii | 58 | 0.0097 |
| East-Asia | Halichoeres trimaculatus | 55 | 0.004 |
| East-Asia | Seriola dumerili | 55 | 0.0042 |
| East-Asia | Nemipterus virgatus | 54 | 0.0003 |
| East-Asia | Plectropomus leopardus | 53 | 0.0076 |
| East-Asia | Epinephelus merra | 53 | 0.0096 |
| East-Asia | Acanthopagrus schlegelii | 49 | 0.0019 |
| East-Asia | Istiophorus platypterus | 48 | 0.004 |
| East-Asia | Acanthurus lineatus | 45 | 0.0065 |
| East-Asia | Naso lituratus | 43 | 0.0059 |
| East-Asia | Euthynnus affinis | 39 | 0.0012 |
| East-Asia | Trimma tevegae | 38 | 0.0008 |
| East-Asia | Upeneus moluccensis | 38 | 0.0056 |
| East-Asia | Scomberoides commersonnianus | 37 | 0.0118 |
| East-Asia | Zanclus cornutus | 36 | 0.0025 |
| East-Asia | Thalassoma lunare | 36 | 0.0045 |
| East-Asia | Macroramphosus scolopax | 36 | 0.0046 |
| East-Asia | Siganus guttatus | 34 | 0.0091 |
| East-Asia | Cephalopholis miniata | 33 | 0.0016 |
| East-Asia | Elagatis bipinnulata | 33 | 0.003 |
| East-Asia | Variola albimarginata | 33 | 0.0055 |
| East-Asia | Diodon holocanthus | 32 | 0.0014 |
| East-Asia | Acanthurus triostegus | 32 | 0.0033 |
| East-Asia | Chaetodon auriga | 32 | 0.0035 |
| East-Asia | Selaroides leptolepis | 32 | 0.0038 |
| East-Asia | Amphiprion sandaracinos | 32 | 0.0044 |
| East-Asia | Ruvettus pretiosus | 32 | 0.0052 |
| East-Asia | Halichoeres claudia | 31 | 0.0053 |
| East-Asia | Thalassoma hardwicke | 30 | 0.007 |
| East-Asia | Decapterus macarellus | 29 | 0.0033 |
| East-Asia | Histrio histrio | 29 | 0.0057 |
| East-Asia | Parupeneus indicus | 29 | 0.0098 |
| East-Asia | Epinephelus malabaricus | 28 | 0.0024 |
| East-Asia | Sicyopterus lagocephalus | 28 | 0.0035 |
| East-Asia | Arothron hispidus | 28 | 0.0051 |
| East-Asia | Priacanthus tayenus | 28 | 0.0089 |
| East-Asia | Sphyraena barracuda | 27 | 0.0008 |
| East-Asia | Anoplogaster cornuta | 27 | 0.0044 |
| East-Asia | Centropyge bispinosa | 27 | 0.006 |
| East-Asia | Leiognathus equulus | 27 | 0.009 |
| East-Asia | Diplospinus multistriatus | 26 | 0.0004 |
| East-Asia | Lutjanus gibbus | 26 | 0.0011 |
| East-Asia | Chaetodon vagabundus | 26 | 0.0015 |
| East-Asia | Pagrus major | 25 | 0.0013 |
| East-Asia | Chaetodon melannotus | 25 | 0.0039 |
| East-Asia | Kuhlia mugil | 25 | 0.0053 |
| East-Asia | Abudefduf sexfasciatus | 25 | 0.0062 |
| East-Asia | Gomphosus varius | 24 | 0.0003 |
| East-Asia | Centropyge vrolikii | 24 | 0.0018 |
| East-Asia | Chlorurus spilurus | 24 | 0.0063 |
| East-Asia | Cookeolus japonicus | 24 | 0.0068 |
| East-Asia | Epinephelus ongus | 24 | 0.0092 |
| East-Asia | Thalassoma quinquevittatum | 24 | 0.0097 |
| East-Asia | Iniistius dea | 23 | 0.0049 |
| East-Asia | Parupeneus multifasciatus | 23 | 0.0054 |
| East-Asia | Epinephelus quoyanus | 23 | 0.0064 |
| East-Asia | Arothron nigropunctatus | 22 | 0.0021 |
| East-Asia | Hemigymnus melapterus | 22 | 0.003 |
| East-Asia | Sciaenops ocellatus | 22 | 0.0032 |
| East-Asia | Acanthurus xanthopterus | 22 | 0.005 |
| East-Asia | Forcipiger flavissimus | 22 | 0.0063 |
| East-Asia | Macropharyngodon meleagris | 21 | 0.0018 |
| East-Asia | Mulloidichthys flavolineatus | 21 | 0.002 |
| East-Asia | Priacanthus macracanthus | 21 | 0.0055 |
| East-Asia | Aluterus monoceros | 20 | 0.0002 |
| East-Asia | Epinephelus stictus | 20 | 0.0008 |
| East-Asia | Grammistes sexlineatus | 20 | 0.0027 |
| East-Asia | Acanthurus nigricauda | 20 | 0.0031 |
| East-Asia | Pristipomoides filamentosus | 20 | 0.0037 |
| East-Asia | Novaculichthys taeniourus | 20 | 0.0052 |
| East-Asia | Zebrasoma scopas | 20 | 0.0062 |
| East-Asia | Scomber australasicus | 20 | 0.0092 |
| East-Asia | Nemateleotris magnifica | 20 | 0.0103 |
| East-Asia | Coris gaimard | 19 | 0.0022 |
| East-Asia | Argyropelecus gigas | 19 | 0.0026 |
| East-Asia | Acanthurus olivaceus | 19 | 0.0029 |
| East-Asia | Thunnus orientalis | 18 | 0.0004 |
| East-Asia | Thalassoma amblycephalum | 18 | 0.001 |
| East-Asia | Diodon hystrix | 18 | 0.002 |
| East-Asia | Selar crumenophthalmus | 18 | 0.0033 |
| East-Asia | Amphiprion polymnus | 18 | 0.0035 |
| East-Asia | Centropyge bicolor | 18 | 0.0037 |
| East-Asia | Nannosalarias nativitatis | 18 | 0.004 |
| East-Asia | Pristigenys niphonia | 18 | 0.0044 |
| East-Asia | Pygoplites diacanthus | 18 | 0.0056 |
| East-Asia | Thalassoma lutescens | 18 | 0.0061 |
| East-Asia | Epibulus insidiator | 18 | 0.0061 |
| East-Asia | Echeneis naucrates | 18 | 0.0081 |
| East-Asia | Abudefduf sordidus | 17 | 0.0023 |
| East-Asia | Awaous grammepomus | 17 | 0.0042 |
| East-Asia | Uranoscopus oligolepis | 17 | 0.0049 |
| East-Asia | Eubleekeria splendens | 17 | 0.0064 |
| East-Asia | Paracanthurus hepatus | 17 | 0.007 |
| East-Asia | Oxycheilinus digramma | 17 | 0.0087 |
| East-Asia | Centropyge tibicen | 16 | 0.0018 |
| East-Asia | Glossogobius aureus | 16 | 0.002 |
| East-Asia | Mulloidichthys vanicolensis | 16 | 0.0028 |
| East-Asia | Lutjanus sebae | 16 | 0.0031 |
| East-Asia | Lutjanus johnii | 16 | 0.008 |
| East-Asia | Mugilogobius chulae | 16 | 0.009 |
| East-Asia | Alepes vari | 15 | 0.0018 |
| East-Asia | Etelis coruscans | 15 | 0.0026 |
| East-Asia | Gerres limbatus | 15 | 0.0028 |
| East-Asia | Stethojulis bandanensis | 15 | 0.0043 |
| East-Asia | Chaetodon lunula | 15 | 0.0044 |
| East-Asia | Apolemichthys trimaculatus | 15 | 0.0057 |
| East-Asia | Synagrops japonicus | 15 | 0.0068 |
| East-Asia | Poromitra crassiceps | 15 | 0.0118 |
| Mediterranean | Scomber scombrus | 362 | 0.0026 |
| Mediterranean | Diplodus vulgaris | 131 | 0.0021 |
| Mediterranean | Trachurus trachurus | 130 | 0.0051 |
| Mediterranean | Boops boops | 69 | 0.0023 |
| Mediterranean | Makaira nigricans | 59 | 0.0041 |
| Mediterranean | Serranus cabrilla | 58 | 0.0093 |
| Mediterranean | Seriola dumerili | 55 | 0.0042 |
| Mediterranean | Mullus barbatus | 53 | 0.0044 |
| Mediterranean | Kajikia albida | 52 | 0.0017 |
| Mediterranean | Mullus surmuletus | 49 | 0.0041 |
| Mediterranean | Istiophorus platypterus | 48 | 0.004 |
| Mediterranean | Trachurus mediterraneus | 47 | 0.0038 |
| Mediterranean | Caranx crysos | 46 | 0.0007 |
| Mediterranean | Pagrus pagrus | 45 | 0.0093 |
| Mediterranean | Balistes capriscus | 44 | 0.0044 |
| Mediterranean | Epinephelus marginatus | 42 | 0.0043 |
| Mediterranean | Serranus hepatus | 38 | 0.0034 |
| Mediterranean | Pagellus acarne | 38 | 0.0011 |
| Mediterranean | Gobius niger | 37 | 0.0099 |
| Mediterranean | Sarpa salpa | 37 | 0.0036 |
| Mediterranean | Macroramphosus scolopax | 36 | 0.0046 |
| Mediterranean | Arnoglossus laterna | 36 | 0.0029 |
| Mediterranean | Spicara smaris | 32 | 0.0059 |
| Mediterranean | Sarda sarda | 31 | 0.0111 |
| Mediterranean | Callionymus lyra | 29 | 0.0008 |
| Mediterranean | Trachurus picturatus | 29 | 0.0037 |
| Mediterranean | Dentex dentex | 28 | 0.004 |
| Mediterranean | Serranus scriba | 27 | 0.0093 |
| Mediterranean | Tripterygion melanurum | 25 | 0.0026 |
| Mediterranean | Pagrus auriga | 25 | 0.0028 |
| Mediterranean | Myctophum punctatum | 24 | 0.0026 |
| Mediterranean | Sparisoma cretense | 23 | 0.0025 |
| Mediterranean | Cepola macrophthalma | 23 | 0.0008 |
| Mediterranean | Sphyraena sphyraena | 23 | 0.0047 |
| Mediterranean | Argyrosomus regius | 23 | 0.0031 |
| Mediterranean | Hygophum benoiti | 21 | 0.0014 |
| Mediterranean | Scorpaena porcus | 21 | 0.0039 |
| Mediterranean | Hoplostethus mediterraneus | 20 | 0.0018 |
| Mediterranean | Trachinus draco | 20 | 0.0028 |
| Mediterranean | Hygophum hygomii | 18 | 0.0062 |
| Mediterranean | Blennius ocellaris | 18 | 0.0049 |
| Mediterranean | Diodon hystrix | 18 | 0.002 |
| Mediterranean | Pagrus caeruleostictus | 17 | 0.0042 |
| Mediterranean | Priacanthus arenatus | 16 | 0.0033 |
| Mediterranean | Pomatoschistus bathi | 15 | 0.0047 |
| Mediterranean | Cyclothone microdon | 15 | 0.0013 |
| Mediterranean | Trichiurus lepturus | 15 | 0.0051 |
